# Monocyte production of C1q potentiates CD8^+^ T cell effector function following respiratory viral infection

**DOI:** 10.1101/2023.06.04.543430

**Authors:** Taylor Eddens, Olivia B. Parks, Dequan Lou, Li Fan, Jorna Sojati, Manda Jo Ramsey, Lori Schmitt, Claudia M. Salgado, Miguel Reyes-Mugica, Tim D. Oury, Craig Byersdorfer, Kong Chen, John V. Williams

**Author notes:** denotes co-first authorship (Taylor Eddens, Olivia B. Parks).

## Abstract

Respiratory viral infections remain a leading cause of morbidity and mortality. Using a murine model of human metapneumovirus (HMPV), we identified recruitment of a C1q-producing inflammatory monocyte population concomitant with viral clearance by adaptive immune cells. Genetic ablation of C1q led to reduced CD8^+^ T cell function. Production of C1q by a myeloid lineage was sufficient to enhance CD8^+^ T cell function. Activated and dividing CD8^+^ T cells expressed a putative C1q receptor, gC1qR. Perturbation of gC1qR signaling led to altered CD8^+^ T cell IFN-γ production and metabolic capacity. Autopsy specimens from fatal respiratory viral infections in children demonstrated diffuse production of C1q by an interstitial population. Humans with severe COVID-19 infection also demonstrated upregulation of gC1qR on activated and rapidly dividing CD8^+^ T cells. Collectively, these studies implicate C1q production from monocytes as a critical regulator of CD8^+^ T cell function following respiratory viral infection.

## Introduction

Lower respiratory tract infections are a leading infectious cause of morbidity and mortality globally, with viral pathogens representing key causative agents.(GBD 2019 LRI Collaborators, 2022; Li and Nair, 2022) This was undoubtedly highlighted by the emergence of SARS-CoV-2 in 2019, which has accounted for over 6 million deaths globally between 2020-20222. In 2022, viruses such as respiratory syncytial virus (RSV), human metapneumovirus (HMPV), and influenza have re-emerged in the setting of waning public health mitigation measures.(Foley et al., 2022; Olsen et al., 2021) Given the constant threat of both new and old respiratory viral pathogens, it is critical to better understand fundamental pathways involved in lung protection and immunopathology.

CD8^+^ antiviral T cells and the humoral compartment are key contributors to acute viral clearance and memory formation.(Martin and Badovinac, 2018; Schmidt and Varga, 2018) Prior animal models have demonstrated formation of protective antigen-specific CD8^+^ T cells in response to viral pathogens, which are subsequently inhibited by a number of mechanisms to limit immunopathology.(Chang and Braciale, 2002; Erickson et al., 2012, 2014; Lawrence et al., 2005; Ruckwardt et al., 2009; Sun et al., 2009) Immunopathology following respiratory viral pathogens has also been attributed to inflammatory monocyte recruitment (Channappanavar et al., 2016; Coates et al., 2018; Nuriev and Johansson, 2019). As more studies have explored the pathogenesis of SARS-CoV-2, potential pathologic contributions of both CD8^+^ T cells and monocytes have been described.(Grant et al., 2021; Knoll et al., 2021; Liao et al., 2020; Merad and Martin, 2020; Singh et al., 2021; Zhou et al., 2020) However, the mechanisms and signaling mediators governing monocyte/CD8^+^ T cell interactions are not fully defined.

In this study, we identify a novel role of C1q-producing inflammatory monocytes during respiratory viral infection. In a murine model of HMPV, C1q was required for optimal CD8^+^ T cell effector function, which express a putative complement receptor gC1qR. In human samples, the C1q/gC1qR axis was present on recruited cells and CD8^+^ T cells, respectively, following severe respiratory viral infection.

## Results

### C1q is produced by an inflammatory macrophage population following human metapneumovirus in mice

Single cell RNA sequencing using the 10x Genomics platform was performed on lung tissue from mock-infected vs HMPV-infected mice on day 7 post-infection, a time in which viral burden is rapidly cleared due to adaptive immunity. An average of 5,730 cells were captured/sequenced per sample, with an average of 24,964 reads per cell. Using clustering analysis, 9 unique cellular populations were identified: T cells, B cells, an inflammatory monocyte, endothelial cells, monocytes, fibroblasts, macrophages, NK cells, and neutrophils (**Fig. 1A**). The inflammatory monocyte population, which clustered closely to a shared monocyte population between HMPV- and mock-infected animals, was only observed following HMPV infection (**Fig. 1B**). HMPV-infected animals also had an expansion of the T cell compartment (**Fig. 1B**). This expansion was partially comprised of CD8^+^ cells that expressed granzyme B (Gzmb), PD-1 (*Pdcd1*) and IFN-γ (*Ifng*) (**Fig. S1A**), consistent with prior reports.(Erickson et al., 2012, 2016) Upregulated ligands and the putative signaling pathways they regulate were modeled using NicheNet (Browaeys et al., 2020). Of the 18 prioritized ligands, *Cd274* (PD-L1) was expressed in several populations and upregulated following HMPV infection (**Fig. S1B**). Further corroborating prior studies, PD-L1 was predicted to regulate several target genes, including *Ifng* (**Fig. S1B**). Collectively, these data demonstrate that the single cell RNA sequencing methodology adequately captured biologic processes consistent with prior work, validating use of this dataset for discovery purposes.

**Figure 1.**
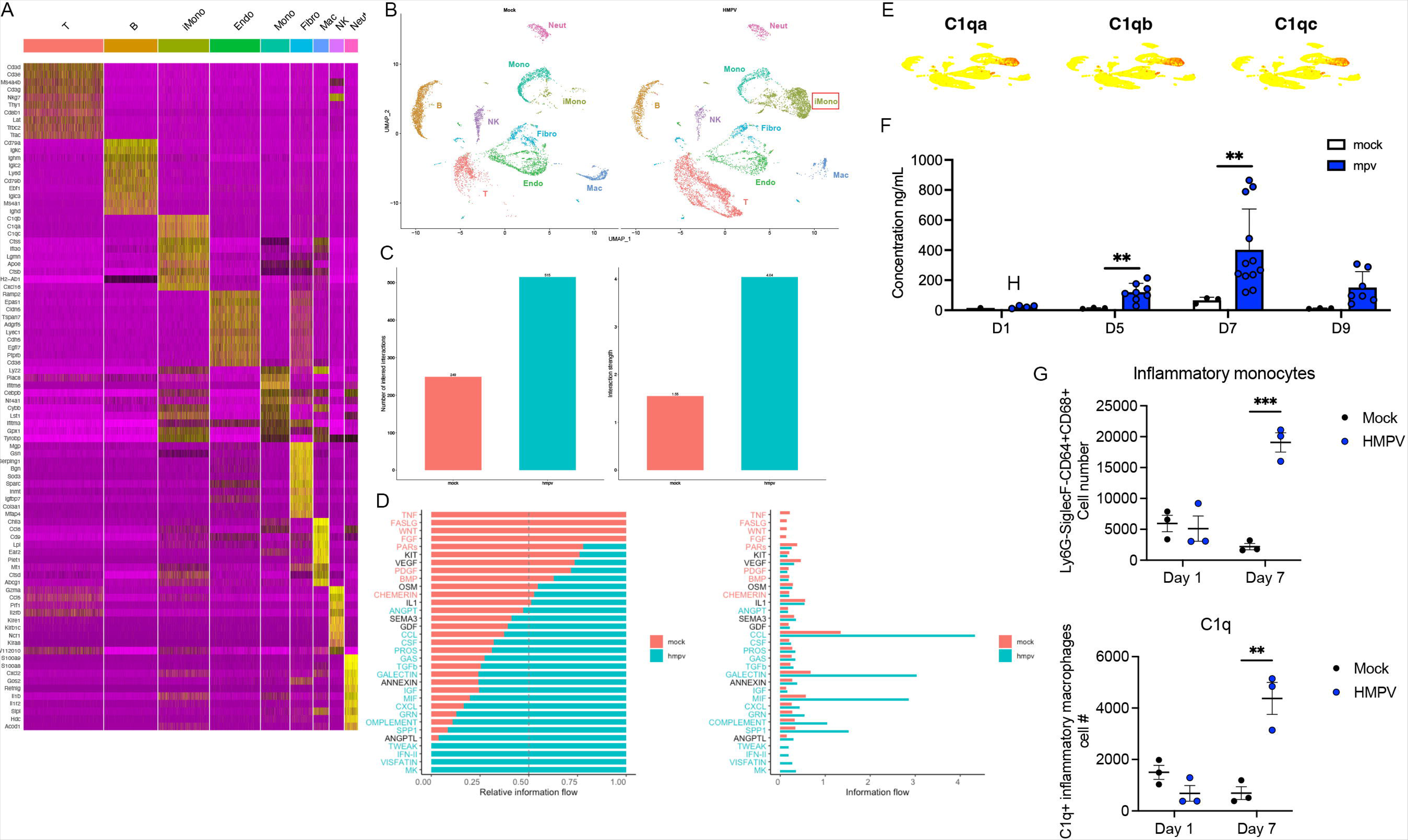
Identification of a C1q signature on day 7 after HMPV infection. A.) Heat map demonstrating clustering of 9 different cell populations after single cell RNA sequencing. B.) Uniform Manifold Approximation and Projection (UMAP) representation of cell populations in mock infected vs. HMPV-infected animals demonstrating an increased inflammatory monocyte (iMono) and T cell population post-infection. C.) CellChat analysis showing number of inferred interations (left) and interaction strength (right) in mock and HMPV infected animals. D.) Pathway analysis of information flow using CellChat. E.) Expression of *C1qa*, *C1qb*, and *C1qc* shown as UMAPs, demonstrating production in iMonos. F.) C1q concentration in bronchoalveolar lavage in mock or HMPV infected animals showing a peak at day 7 post infection. *p<0.01 by multiple t-tests. G.) Quantification of inflammatory monocytes (top) and C1q-producing iMonos (bottom) on day 1 and 7 post-exposure showing increased populations at day 7. *p<0.01, *** p<0.005 by one-way ANOVA with multiple comparisons.

We next sought to analyze cell-to-cell communication in an unbiased manner using Cellchat (Jin et al., 2021). HMPV-infected animals had upregulation of both interaction number and strength (**Fig. 1C**). Cellchat pathway analysis revealed a strong complement signature in HMPV-infected animals (**Fig. 1D**). Inflammatory monocytes were the largest contributors of outgoing signals in number and strength, while T cells were the largest node of signals received (**Fig. S2A-B**). Inflammatory monocytes had several methods of communicating to T cells when analyzed via Cellchat, including through chemotactic mechanisms (**Fig. S2C**). However, analyzing the transcripts within inflammatory monocytes, *C1qa*, *C1qb*, and *C1qc* had the highest expression (**Fig. 1E**). In fact, upon analyzing globally differentially expressed genes in HMPV-infected animals compared to mock infected animals, *C1qa*, *C1qb*, and *C1qc* were the three highest upregulated genes in the lung (*p*<1×10^−50^).

C1q was also upregulated at the protein level in bronchoalveolar lavage fluid over the course of HMPV infection, peaking at day 7 post-infection (**Fig. 1F**). Additionally, using validated surface markers for various monocyte/macrophage populations, we next identified the inflammatory monocyte population via flow cytometry (**Fig. 1G**, **Fig. S3A-B**). Notably, there was a significant increase in C1q-expressing inflammatory monocytes at day 7 post-HMPV infection (**Fig. 1G, Fig. S3C**). C1q production was not seen in other myeloid cells, such as neutrophils (**Fig S3D**). Additionally, if staining was performed prior to permeabilization, the C1q signal in the inflammatory monocyte population disappeared, suggesting intracellular production rather than detection of bound C1q (**Fig S3E**). Collectively, these data suggest HMPV-infected animals have a robust C1q-producing inflammatory monocyte population present late in HMPV infection, coincident with adaptive immunity.

### C1q is required for optimal CD8 effector function

We next assessed CD8^+^ T cell responses in the absence of C1q using 6(Cg)*C1qa^tm1d(EUCOMM)Wtsi^*/TennJ (referred to as *C1qa*^−/−^) mice following infection with HMPV (TN/94-49), a well characterized strain that causes mild disease in adult B6 mice.(Erickson et al., 2012) Lack of C1q induction was confirmed in *C1qa^−/−^* mice via ELISA (**Fig. S4A**). There was no difference in weight loss or viral titer between *C1qa*^−/−^ and B6 mice (**Fig. S4B-C**). At day 7 post-infection, there was also no difference in frequency or absolute cell number of epitope-specific CD8^+^ T cells using an MHC-I tetramer bearing one of the immunodominant H2-K^b^ HMPV epitopes, N_11-18_(**Fig. 2A**). Similarly, inhibitory receptors (i.e. PD-1, TIM-3, LAG-3, and 2B4), activation states (CD44, CD62L), and transcription factors (i.e. T-bet, GATA3, Foxp3, EOMES, TOX, and TCF1) on virus-specific CD8^+^ T cells were unchanged between the two groups (**Fig. S4D-F**). B6 and *C1qa^−/−^* mice displayed a similar extent of immunopathology following TN/94-49 infection. **(Fig. S4G-H).** However, CD8^+^ T cells from *C1qa*^−/−^ mice exhibited a striking decrease in function, producing significantly less granzyme B, IL-2, and IFN*γ* (**Fig. 2B**). Combinatorial analysis assessing CD107a, IFN*γ*, granzyme B, IL-2, and perforin production revealed that *C1qa*^−/−^ CD8^+^ T cells were less polyfunctional compared to CD8^+^ T cells from B6 mice (**Fig. 2C**). To determine the effects of C1q signaling to CD8^+^ T cells during severe HMPV infection, we used a clinical isolate of HMPV, C-202, known to cause severe disease in young adult mice (manuscript submitted). *C1qa*^−/−^ mice were able to clear the virus, but lost significantly more weight (**Fig. S5A**) and had higher clinical scores compared to B6 mice (**Fig. S5B**). *C1qa^−/−^* mice also enhanced histopathology following C-202 infection when compared to B6 mice (**Fig. S5C-D**). These data indicate that the absence of C1q signaling has detrimental effects during severe respiratory disease.

**Figure 2.**
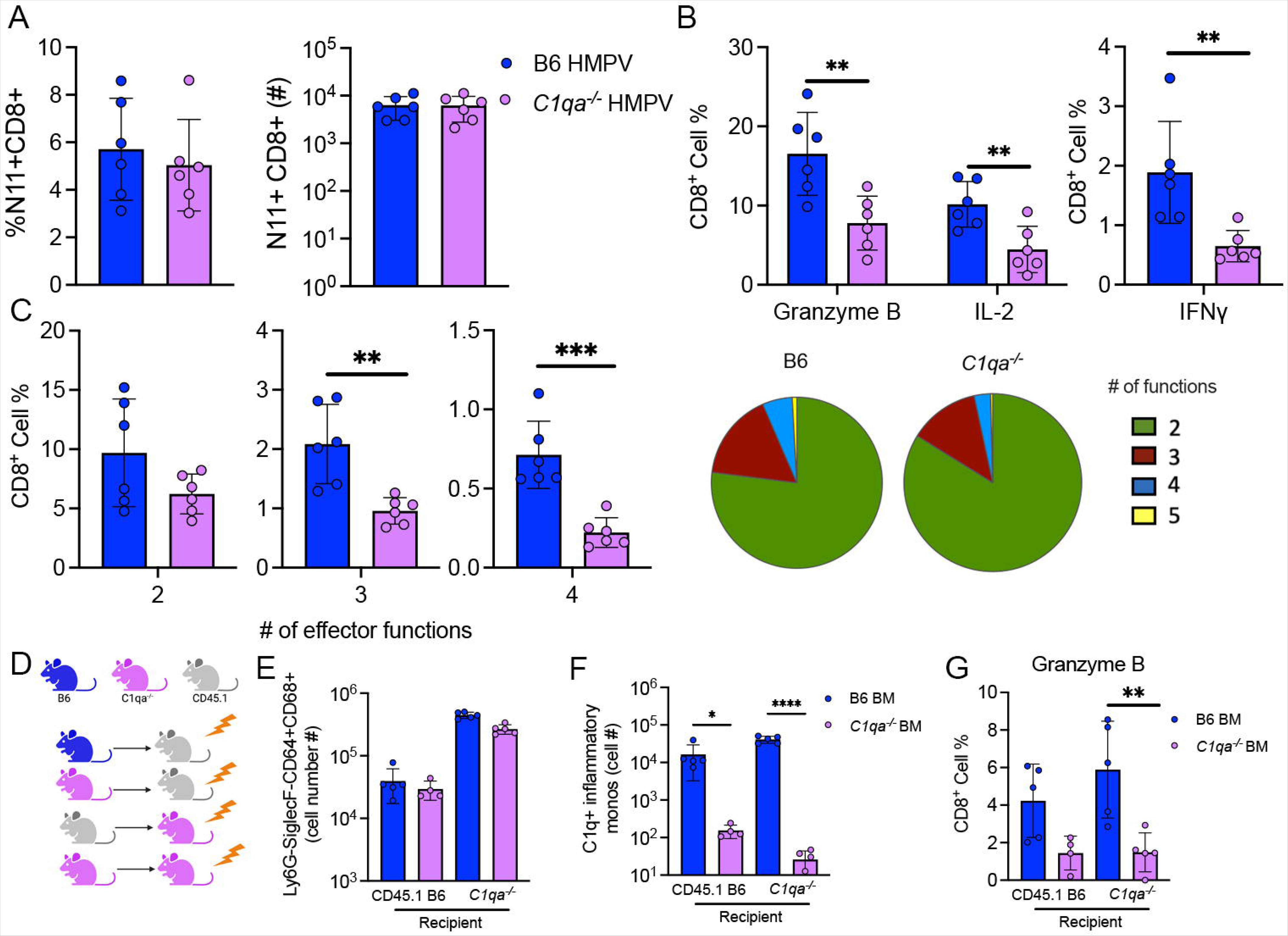
Absence of C1q leads to less functional CD8^+^ T cells. A.) There was no difference in N11^+^ CD8^+^ T cells between B6 and *C1qa^−/−^* in cell percent and absolute cell number. B.) CD8^+^ T cells in *C1qa^−/−^* mice had impaired production of effector cytokines, granzyme B, IL-2, and IFN-γ. C.) Combinatorial analysis of functional markers revealed that CD8 T cells from *C1q* mice produced significantly less 3 or 4 markers. Bar graphs (left) show raw data, pie charts (right) reflect summary of raw data. D.) Experimental schematic for four-way transplant. E.) There was no difference in absolute cell number of inflammatory monocytes between the four transplant groups. F.) The recipients that received B6 BM had significantly more C1q producing inflammatory monocytes compared to recipients that received *C1qa^−/−^* BM. G.) Recipient mice that received *C1qa^−/−^* BM had CD8^+^ T cells that produced less granzyme B compared to mice that received B6 BM. *p<0.05, ** p<0.01, *** p<0.005, **** p<0.001 by unpaired t-test or one-way ANOVA with multiple comparisons.

To determine if this CD8^+^ T cell phenotype required the complement pathway downstream of C1q, we used *C3^−/−^*which lack C3, a common protein utilized by all three complement pathways **(Fig. S6A).** There was no difference in weight loss **(Fig. S6B).** In contrast to WT B6 mice, *C3^−/−^* mice had detectable titer **(Fig. S6C)** at day post-infection. However, there were no differences in epitope-specific CD8^+^ T cell frequency in *C3^−/−^* mice **(Fig. S6D).** There was also no difference in inhibitory receptor **(Fig. S6E),** transcription factor expression **(Fig. S6E),** or polyfunctionality of CD8^+^ T cells **(Fig. S6F-G)** between *C3^−/−^* and B6 mice, suggesting the alternative complement activation and shared membrane attack complex formation pathway is not involved in the CD8 disfunction.

To assess if myeloid-derived C1q was required for enhanced CD8^+^ T cell function, we utilized a four-way reciprocal bone marrow (BM) chimera approach. First, *C1qa*^−/−^ or B6 BM was transplanted into lethally irradiated CD45.1 recipients or the reciprocal where CD45.1 BM was transplanted into lethally irradiated *C1qa*^−/−^ and B6 recipients (**Fig. 2D**). For clarity, the experimental groups are referred to as: BM → host (i.e. *C1qa*^−/−^ BM into CD45.1 recipient = *C1qa*^−/−^ → CD45.1) There was no difference in viral titer between transplant groups. There were equal numbers of inflammatory monocytes in *C1qa*^−/−^ → CD45.1 and B6 → CD45.1 groups (**Fig. 2E**), but *C1qa*^−/−^ → CD45.1 had negligible C1q^+^ inflammatory macrophages compared to B6_BM_ → CD45.1_H_ (**Fig. 2F**). Compared to B6_BM_ → CD45.1_H_, CD8^+^ T cells from the *C1qa*^−/−^ → CD45.1 group produced significantly less granzyme B (**Fig. 2G**), while minimal antigen-specific IL-2/IFN-γ was noted in either group (**Fig. S7A**).

To demonstrate *C1qa^−/−^* CD8^+^ T cells were not intrinsically defective, a competitive adoptive transfer experiment of congenically labeled CD45.1 B6 vs CD45.2 *C1qa^−/−^* CD8^+^ T cells into the same *Rag1^−/−^* recipient with reconstituted with CD45.1 CD4^+^ T and B lymphocytes was performed (**Fig. S7B**). No difference was observed in granzyme B or IFN-γ production in B6 vs *C1qa^−/−^* CD8^+^ T cells in this model (**Fig. S7C**).

### Blockade of gC1qR leads to reduced CD8 effector function

As mice lacking myeloid-derived C1q had reduced CD8^+^ T cell function following HMPV infection, we next assessed if C1q was directly acting on CD8^+^ T cells. Recombinant C1q robustly bound to purified lung CD8^+^ T cells from day 7 post-HMPV infected *C1qa^−/−^* animals (**Fig. 3A**). One putative receptor for C1q, gC1qR (aka C1QBP), was expressed on a subset of CD8^+^ T cells in the prior single cell analysis (**Fig. 3B**).(Ghebrehiwet et al., 2019) Moreover, further analysis of transcriptional profiles of CD8^+^ T cells revealed that gC1qR expression was confined to proliferating, cytotoxic CD8^+^ T cells with increased expression of *Mik67* and *Gzmb* (**Fig. 3B**). After identification of gC1qR as a possible receptor for C1q on CD8^+^ T cells, we subsequently utilized a blocking antibody on *ex vivo* stimulated lung cells from mice infected with HMPV. Production of IFN-γ was abrogated in cells treated with N11 (a dominant MHC class I peptide) and IZl-gC1qR antibody (**Fig. 3C**). As described previously, IFN-γ production was enhanced following blockade of PD-L1 (**Fig. 3C**).(Erickson et al., 2014) However, IZlgC1qR treatment reduced IFN-γ production even in the presence of PD-L1 blockade, suggesting two distinct pathways of IFN-γ regulation in CD8^+^ T cells (**Fig. 3C**). IL-2, IFN-γ, and downstream IFN response genes CXCL10 and CXC9 protein levels were also decreased when cells were treated with anti-gC1qR and combination treatment; again, C1qR blockade reduced production of these cytokines despite the presence of PD-L1 blockade (**Fig. S8A**). Moreover, cells treated with recombinant C1q also demonstrated an upregulation of IFN-γ production, which was again blunted by IZlgC1qR treatment (**Fig. 3D**). C1q has previously been shown to regulate CD8^+^ T cell metabolism and function in a chronic viral infection model.(Ling et al., 2018) To that end, purified CD8^+^ T cells isolated from the lungs at day 7 post-HMPV infection showed a diminished spare respiratory capacity (SRC) when treated with IZlgC1qR antibody (**Fig. 3E**). Collectively, these data demonstrate a critical role for gC1qR receptor in maintaining optimal CD8^+^ T cell effector function and metabolic capacity that functions independently of PD-L1 signaling.

**Figure 3.**
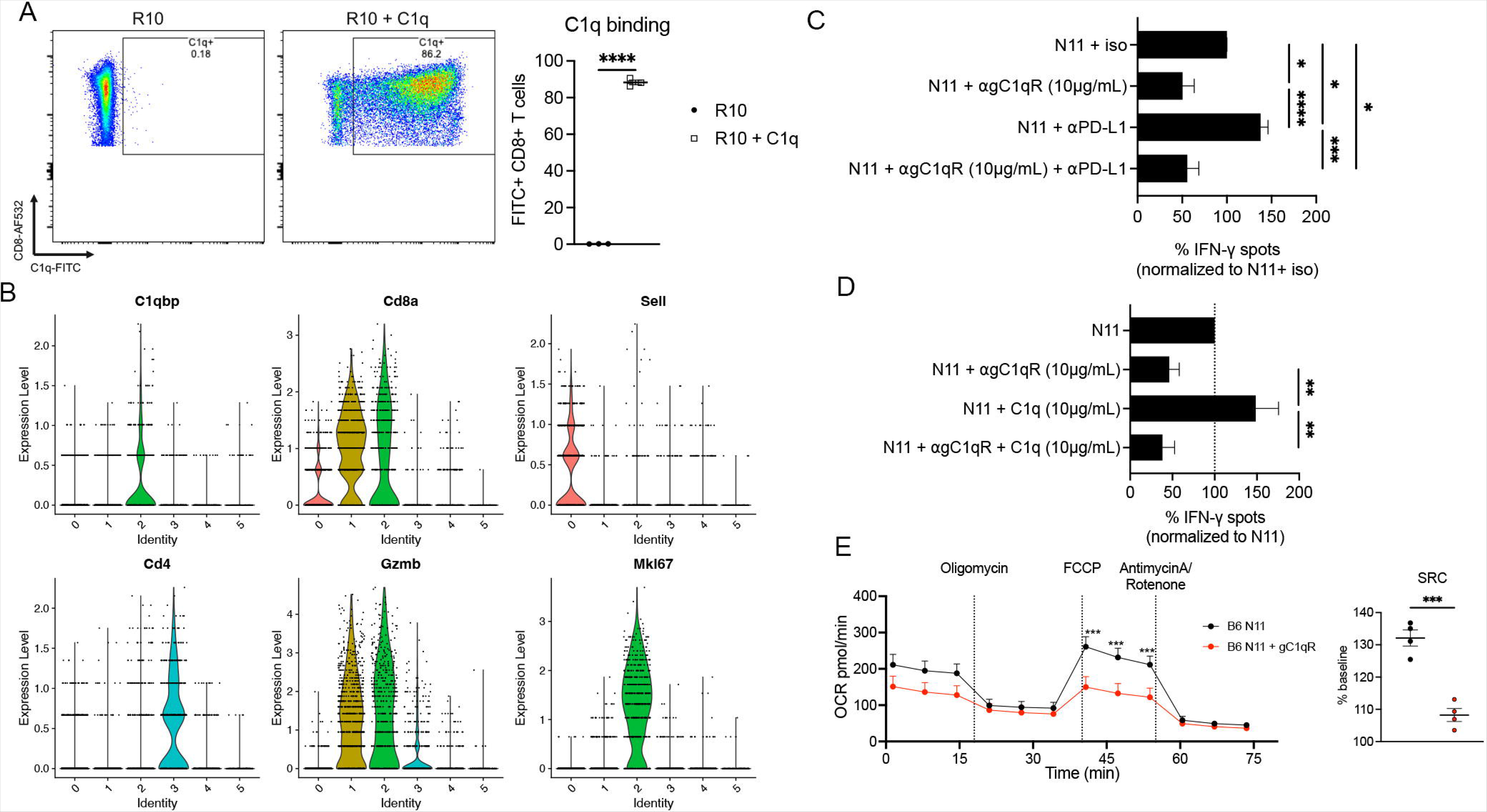
gC1qR blockade leads to reduced CD8^+^ T cell function. A.) C1q bound to cultured murine CD8^+^ T cells. Representative flow plots (left), quantification of C1q-FITC^+^ CD8^+^ T cells (right). B.) Violin plots showing expression of *C1QBP* (i.e gC1qR), *CD8*, *SELL*, *CD4*, *GZMB*, and *MIK67* from a prior single cell analysis (Ghebrehiwet et al., 2019). C.) IFN-γ was significantly decreased in murine lung lymphocytes undergoing *ex vivo* Class I peptide stimulation on day 7 p.i. when αgC1qR was added. While PDL1 blockade increased IFN-γ production. Combination treatment resulted in decreased IFN-γ production. D.) Addition of recombinant C1q increased IFN-γ production. E.) Purified CD8^+^ T cells on day 7 p.i. had diminished spare respiratory capacity (SRC) when treated with αgC1qR. *p<0.05, ** p<0.01, *** p<0.005, **** p<0.001 by unpaired t-test or one-way ANOVA with multiple comparisons.

### C1q in human respiratory viral infection

The myeloid populations capable of producing C1q is one of myriad differences between murine and human immunology. Human alveolar macrophages produce C1q at steady state, while C1q production in mice is often limited to interstitial macrophages and recruited alveolar macrophages post-inflammatory stimulus (Aegerter et al., 2022; Leach et al., 2020; Li et al., 2022). However, we tested whether respiratory viral infection altered C1q production in the lung by using immunofluorescent staining on archived pediatric lung specimens. In a child who had a partial lung resection due to pleural blebs, C1q was visualized in the alveolar spaces (**Fig. 4A, top**). In a case of fatal HMPV, the location of C1q+ cells markedly shifted to the interstitial and perivascular spaces (**Fig. 4A, middle**). Likewise, staining from a fatal case of mixed rhinovirus/parainfluenza virus pneumonia showed numerous interstitial and perivascular inflammatory cells producing C1q (**Fig. 4A, bottom**).

**Figure 4.**
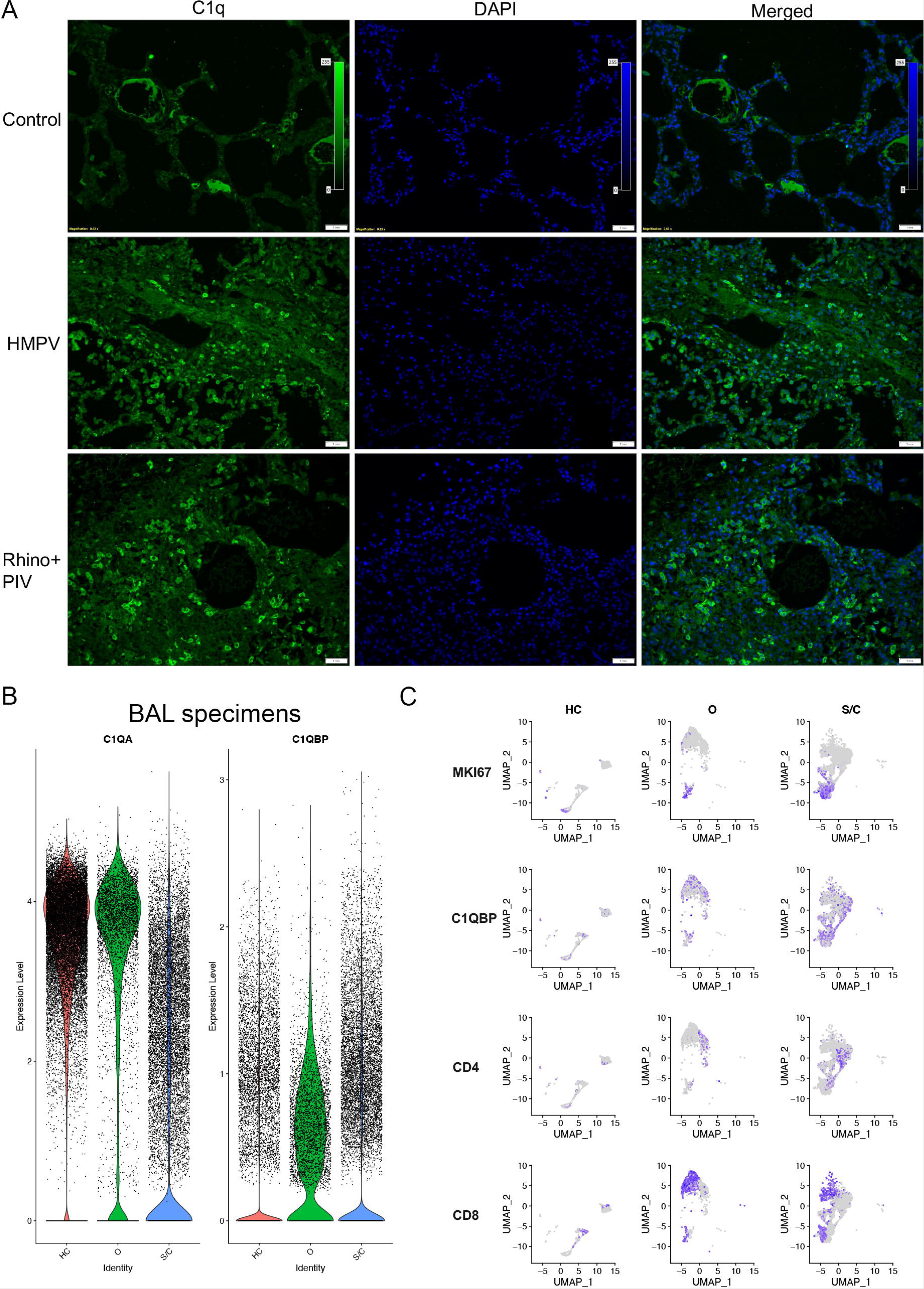
Evidence of C1q:gC1qR axis in humans with respiratory viral infections. A.) Immunofluorescent staining of human lung tissue (top: resection from child with pleural blebs, middle: autopsy specimen from a child who succumbed to HMPV, bottom: autopsy specimen from a child who succumbed to rhinovirus/parainfluenza (PIV) infection). C1q (green) is present in alveolar spaces in healthy child with re-distribution to interstitial spaces in infectious conditions. B.) *C1QA* expression (left) is reduced in severe COVID-19 (S/C) infection compared to healthy controls (HC) or moderate cases (O). *C1QBP* expression (right) is detectable in BAL specimens. C.) UMAPs of *MKI67* (marker of rapid division), *C1QBP, CD4,* and *CD8* in healthy contorls, moderate COVID, and severe COVID cases demonstrating upregulation of *C1QBP* in a subset of rapidly dividing CD8s in severe COVID.

We next queried publicly available single-cell RNA sequencing data from bronchoalveolar immune cells in humans with COVID-19 infection.(Liao et al., 2020) Healthy control participants and moderate COVID-19 patients had high levels of C1q production at baseline, while severe COVID-19 patients had a loss of C1q expression (**Fig. 4B**). Evaluation of the T cell compartment in this data set demonstrated an increase in both the CD8^+^ and CD4^+^ T cell populations in moderate and severe COVID-19 disease (**Fig. 4C**). Notably, gC1qR (gene:*C1QBP*) expression could be noted on the CD8^+^ T cell population in moderate COVID-19 disease, but was more abundantly expressed on *MKI67*^+^ CD8^+^ T cells in severe disease (**Fig. 4C**). Collectively, these data demonstrate altered C1q production in the human lung following respiratory viral infection and the presence of a known receptor for globular C1q on CD8^+^ T cells.

## Discussion

Taken together, our data identifies an integral role of C1q signaling in respiratory viral infection. Three key findings from this study include: (1) there is a novel inflammatory monocyte population that produces C1q concomitant with adaptive immune-mediated clearance of virus, (2) C1q directly impacts CD8^+^ T cell function in our murine model and (3) the C1q/gC1qR axis presents a possible mechanism as to how monocyte-derived C1q regulates CD8^+^ T cell function in both mice and humans across multiple respiratory viral pathogens.

C1q is canonically thought of as the initiator of the classical complement pathway in the innate immune response.(Kishore and Reid, 2000) However, C1q has also been implicated in both homeostasis and inflammatory conditions, including clearance of apoptotic debris and recognition of damage/pathogen associated molecular patterns with resultant phagocytosis of extracellular pathogens.(Son, 2022) Previous studies in cancer and autoimmunity models in mice identified inflammatory monocytes and macrophages as key producers of C1q, which facilitates a unique role of C1q independent of the complement cascade.(Bulla et al., 2016; Ling et al., 2018; Ogawa et al., 2022; Roumenina et al., 2019)

Here, we demonstrate a novel role of a C1q-producing myeloid population that promotes CD8^+^ T cell function following respiratory viral infection in our murine model. During HMPV infection, the absence of C1q resulted in diminished effector CD8^+^ T cell polyfunctionality and severe clinical disease when infected with a virulent strain of HMPV. In contrast to our data, in chronic LCMV infection in *C1qa^−/−^* mice, CD8^+^ T cells differentiated into short-lived effector cells with diminished metabolic capacity but more potent effector activity.(Ling et al., 2018) These data suggest that the duration of infection/exposure to antigen may play a role in this regulatory pathway on CD8^+^ T cells, as pathogen-specific antibody and/or immune complex formation may alter the effects of C1q on CD8^+^ T cells.

There are differences between murine and human C1q production in the lung. In the human lung, alveolar macrophages produce C1q at baseline, while mice have negligible C1q production in the absence of infection.(Aegerter et al., 2022; Leach et al., 2020; Li et al., 2022) In staining of human lung tissue, C1q could be distinctly seen in the alveolar spaces of a lung section from an uninfected child; this distribution changed dramatically to an interstitial population in the setting of lethal respiratory viral infection, consistent with our animal model. Additionally, single cell sequencing of the myeloid compartment in humans infected with COVID-19 demonstrated C1q as a differentially expressed gene in alveolar macrophages, although some expression was observed in monocyte-derived macrophages.(Melms et al., 2021) Interestingly, C1q transcript was lost in the cells isolated from bronchoalveolar lavage fluid from patients with severe COVID; these data, coupled with the immunofluorescent images shown in our study, may suggest the cell type and location of C1q production shifts during acute respiratory viral infection in humans.

The next question we addressed is how might communication between C1q and CD8^+^ T cells be occurring. C1q has been previously shown to bind to human CD8^+^ T cells, with increased binding following *in vitro* stimulation (Chen et al., 1994). Similarly, our data shows that murine CD8^+^ T cells following HMPV infection readily bound C1q. While there are several putative receptors for C1q, gC1qR (i.e. C1q binding protein, C1qBP) recognizes the globular head of the C1q polypeptide and plays a critical role in mitochondrial fitness.(Ghebrehiwet et al., 2019; Thielens et al., 2017; Tian et al., 2022) Rapidly dividing CD8^+^ T cells during HMPV infection upregulated gC1qR expression in our murine model and in patients with severe COVID-19. These findings are consistent with a prior study which demonstrated gC1qR is abundantly expressed on human CD8^+^ T cells in the setting of chronic hepatitis C infection.(Yao et al., 2004) Subsequent *in vitro* blockade of gC1qR led to reduced effector function in our model. Blocking gC1qR signaling in CD8^+^ T cells from the murine lung following HMPV infection also led to reduced metabolic capacity, similar to a prior study assessing the metabolomics of *C1qa^−/−^* CD8^+^ T cells in the LCMV chronic model.(Ling et al., 2018) Collectively, these data identify gC1qR as at least one of the critical C1q receptors required for optimal CD8^+^ T cell function in multiple respiratory viral models.

There are limitations and possible future directions from the current study. First, *C1qa^−/−^* mice cleared HMPV comparably to wild type mice, despite the diminished CD8^+^ T cell effector function. However, in our C-202 model and in prior LCMV models, *C1qa^−/−^* mice have increased pathology.(Ling et al., 2018) These findings may suggest effective, but pathologic, alternative mechanisms of clearance in the absence of C1q, which would warrant further study of other anti-viral cell types such as NK cells. Second, restoration of CD8^+^ T cell functionality following B6 hematopoietic stem cell transplantation was noted with granzyme B production, but not other effector functions. These data support C1q’s role in CD8^+^ T cell responses, but may be limited by the negative effects of whole body radiation in the absence of C1q.(Markarian et al., 2021) The interplay between C1q, CD8^+^ T cell function, and pathogen-specific antibody could also be explored mechanistically by passive convalescent serum transfer in *C1qa^−/−^* mice and/or re-challenge experiments. The latter would also be fascinating in the context of CD8^+^ T cell memory formation in the absence of C1q. Third, further evaluation of the C1q/gC1qR axis in both our mouse model and human samples are warranted. gC1qR loss of function is poorly tolerated in T cells, with an *in vitro* knockdown approach in mice resulting in diminished proliferative capacity, increased mitochondrial membrane permeability, and ultimately increased apoptotic cell death.(Tian et al., 2022) This makes it technically challenging to completely knock out the gC1qR receptor on CD8^+^ T cells as this will have detrimental effects to the survival of CD8^+^ T cells, but also underscores the importance of this receptor in CD8^+^ T cell fitness. In a chimeric antigen receptor (CAR) T cell model, heterozygosity of gC1qR lead to reduced CD8^+^ T cell effector function and increased tumor burden (Tian et al., 2022), which supports our hypothesis that gC1qR signaling on CD8^+^ T cells is crucial to their function. One future study could assess serum C1q levels in acute disease or *ex vivo* manipulation of gC1qR from cells isolated from infected patients which could provide insights into the potential diagnostic and/or therapeutic potential of this pathway.

Collectively, the current study identifies a novel C1q-producing monocyte population and elucidates a potential mechanism of how C1q signaling via gC1qR on CD8^+^ T cells can regulate optimal CD8^+^ cytotoxic antiviral effector function during respiratory viral infections.

## Acknowledgements

Supported by NIH AI085062 (JVW), 1F30HL159915 (OBP), T32 GM-008208 (OBP, JS), K12 HD000850 (TE), and the Henry L. Hillman Foundation (JVW). Supported in part by the Marjory K. Harmer endowment for research in Pediatric Pathology (MRM). We thank the University of Pittsburgh Unified Flow Core for help with flow cytometry. We thank the NIH Tetramer Core Facility (contract number 75N93020D00005) for providing HMPV tetramers.

## STAR Methods

### Resource Availability

#### Lead contact

Further information and request for resources and reagents should be directed to and will be fulfilled by the lead contact, John Williams.

#### Materials availability

No new reagents or mouse lines were generated in the study.

#### Data and code availability

Single cell RNA sequencing data generated by this study have been deposited to the GEO database and will be available publicly at the date of publication (accession number: pending). Any additional information required to reanalyze the data reported in this paper is available from the lead contact upon request.

### Experimental model and study participant details

#### Mice and virus stocks

C57BL/6 (strain 664), 6(Cg)-*C1qa^tm1d(EUCOMM)Wtsi^*/TennJ (strain 31675, referred to as *C1qa*^−/−^), B6.129S4-*C3^tm1Crr^*/J (strain 29661, referred to as *C3^−/−^*), *Rag1^−/−^* (strain 2216) and B6.SJL-*Ptprc^a^ Pepc^b^*/BoyJ (strain 2014, referred to as CD45.1) mice were purchased from the Jackson Laboratory. Six-to-eight-week mice were anesthetized with 3% isoflurane and infected with 2.0×10^6^ PFU of HMPV strain TN/94-49 (genotype A2) in 100µL sterile PBS. HMPV was grown in LLC-MK2 cells and purified as previously described.(Williams et al., 2005) Mice were treated with a mock-infected LLC-MK2 lysate as a negative control. In select experiments, mice were infected with 1.0×10^5^ PFU of clinical isolate C2-202 (genotype B1), which is a highly virulent strain of HMPV (manuscript submitted). All animals were handled according to protocols approved by the University of Pittsburgh Institutional Animal Care and Use Committee.

#### Human lung tissue staining

Lung samples from children who died from respiratory viral infections (autopsy cases) were selected on retrospective review of specimens from the Department of Pathology, UPMC Children’s Hospital of Pittsburgh. Normal lung tissues resected as perilesional lung in patients with pleural blebs were used as normal control. Use of cadaveric tissue was approved by the University of Pittsburgh Committee for Oversight of Research and Clinical Training Involving Decedents (CORID).

### Methods

#### Single cell RNA sequencing

The lungs were removed from HMPV-infected or mock-infected mice on day 7 post-infection. The lungs were then minced via scissors, digested with DNAse/collagenase at 37°C for 1 hour, passed through a 70µm strainer, and treated with RBC lysis buffer (ACK, Gibco, Cat:A1049210) to generate a single cell suspension. Cells were then stained with cell hashing antibody from Biolegend following the CITE-seq protocol (https://cite-seq.com/protocols/). After the final wash, cells were passed through a 40uM cell strainer and enumerated by Cellometer2000 before loading onto 10x Chromium controller for cell capture using the 5’ V2 kits. Libraries for gene expression and hash tag oligos were constructed following protocols from 10x Genomics and the New York Genome Center. Final libraries were QCed by Agilent TapeStation then sequenced on an Illumina Novaseq 6000 targeting 50,000 reads per cells. Sequencing data were processed with Cellranger7.0 before downstream analysis using Seurat.

#### Single Cell RNA-Seq data processing

Single cell RNA sequencing analysis was performed using Seurat 4.0 with R (version 4.1.1). Samples were demultiplexed by hash-tag-oligos (HTOs) and doublets identified by HTOs were removed before downstream analysis. Poor-quality droplets were excluded from subsequent analysis if deficient number of genes or high percentage of mitochondrial genes were detected. Differential gene expression was performed using non-parametric Wilcoxon rank sum test. The results were adjusted for multiple comparisons using Bonferroni correction. For human single cell RNA sequencing analysis, the following publicly available dataset was mined: GSE145926.(Liao et al., 2020)

#### C1q ELISA

Quantification of C1q on lung homogenate and bronchoalveolar lavage fluid (BAL) was performed via ELISA following manufacturer’s instructions (Abcam, cat# 291069).

#### Viral burden titration

Viral titers were measured via plaque titration as described previously.(Williams et al., 2005; Xu et al., 2018) Briefly, lung homogenates were serially diluted 1:10 in 0% Opti-MEM media containing 1:2000 trypsin and adsorbed to a monolayer of LLC-MK2 cells in a 24-well tissue culture dish for one hour at room temperature. Methylcellulose-containing overlay media with 1:500 trypsin was added, followed by a 5-day incubation at 37°C. Plates were then fixed with formalin, blocked with PBST containing 5% nonfat dried milk, stained using 1:1000 guinea pig anti-HMPV primary antibody and 1:1000 anti-guinea pig-HRP secondary antibody, and developed using TrueBlue^TM^ substrate (KPL, Cat #5510-0050).

#### Histology

Following dissection and isolation, the lower half of the left lung lobe was insufflated with 10% formalin, embedded in paraffin, sectioned, and stained with H&E. Histologic scoring was performed by a trained pathologist in a blinded fashion.

#### Flow cytometry

Following generation of a single cell suspension from lung as above, cells were then plated for *ex vivo* peptide stimulation or tetramer staining in parallel, as described previously.(Erickson et al., 2012) For peptide stimulation, cells were stimulated with 10µM N11 peptide or irrelevant control peptide in the presence of BFA/monensin and CD107-PE antibody for 5 hours at 37°C. Cells were then stained with live/dead violet (1:1000 in PBS, Invitrogen, Cat:L34964A), washed x2 in FACS, treated with Fc blockade (1:100 in FACS buffer, Tonbo™, Cat:70-0161-M001), and stained for surface markers for 45 mins at 4°C (1.5µL antibody/sample in BD Horizon™ Buffer, cat:566349, Supplemental Table 1). Cells were then fixed with FOXP3 fix/permeabilization buffer for 20 minutes (Invitrogen, Cat:50-112-8857), washed x1 in permeabilization buffer, and stained for intracellular markers for 30 minutes at 4°C(4.5µL antibody/sample in 1:1 mixture of BD buffer and fix/perm buffer, Supplemental Table 1). Cells were resuspended in FACS buffer. For tetramer staining, cells were incubated for 30mins at RT in FACS buffer containing dasatinib, followed by 1:100 N11 class I tetramer or an irrelevant tetramer for 90 minutes at RT in the dark. After live/dead, Fc block, and surface staining as above, cells were fixed for 18 hours with FOXP3 fix/permeabilization buffer. Cells were then washed in fix/perm buffer x2 and resuspended in FACS buffer. Cells were then stained for transcription factors (2.5µL antibody/sample in 1:1 mixture of BD buffer and fix/perm buffer, Supplemental Table 1).

Following two washes, the cells were resuspended in FACS buffer with 100µL of Biolegend Counting Beads. Fluorescence minus one (FMO) controls were used for all inhibitory receptor and transcription factor gating.

For macrophage staining, cells were stained for live/dead, Fc blockade, and surface markers as above (Supplemental Table 1). Cells were then fixed in FOXP3 fix/perm buffer for 20 minutes, stained with 1.5µL anti-C1q-FITC for 1 hour at 4°C, washed, and subsequently resuspended in FACS buffer.

For all conditions, samples were strained through nylon filters and run on a Cytek^®^ Aurora multispectral flow cytometer. Unstained cells from each experiment were generated using 2% PFA and used to minimize autofluorescence. Data analysis was performed with Flowjo(v10.8.1).

#### Bone marrow transplantation and adoptive transfer models

For bone marrow transplantation, CD45.1 or *C1qa^−/−^* recipients were irradiated with two doses of 5.5 Gy 5 hours apart (MultiRad 350, Precision X-Ray Irradiation). Twenty-four hours later, bone marrow from either CD45.2 B6 or *C1qa^−/−^* mice was isolated by flushing the femur/tibia with D-10 media. Irradiated mice then received 2.0×10^6^ cells in 200µL sterile PBS via tail vein injection. Following a 6-week engraftment period, mice were infected with HMPV and analyzed for CD8 function as above.

For adoptive transfer, CD19^+^ cells, CD8^+^ T cells, and CD4^+^ T cells were purified from donor lymph nodes/spleen via magnetic separation. Briefly, CD45.1 B6, CD45.2 B6 or *C1qa^−/−^* lymph nodes and spleens were processed sequentially through 70µm and 40µm filters. For CD4^+^ and CD8^+^ negative selection, cells were incubated for 5 minutes with selection antibody cocktail (Miltenyi Biotec, cat # 130-104-454, 130-104-075) followed by a 10 minute incubation with anti-biotin magnetic beads (cat# 130-090-485). CD19^+^ cells were isolated via positive selection by incubating cells with anti-CD19 biotin conjugated antibody (Invitrogen cat # 13-0193-82), followed by incubation with anti-biotin beads. Labeled cells were then applied to primed LS columns (Miltenyi Biotec cat # 130-042-401) and collected following two washes. Cells were then washed with PBS and counted on a BD Accuri^TM^ cytometer. A mixture of 2.0×10^6^ B6 CD19^+^, 2.0×10^6^ B6 CD4^+^, and a 1:1 mixture CD45.1:*C1qa^−/−^* of 2.0×10^6^ total CD8^+^ T cells was then administered to *Rag1^−/−^* recipients via tail vein injection. Following a 3-week reconstitution period, mice were then infected with HMPV and analyzed as above.

#### IFN-γ ELISpot

Following generation of a single cell suspension from lung, 50,000 cells were plated per well in triplicate on an IFN-γ ELISpot plate (R&D systems, cat# EL485), with 10µM of irrelevant (LCMV gp66-77 peptide) or a MHC class I restricted immunodominant HMPV epitope (N11). Antibody blockade was performed with 10µg/mL anti-gC1qR (clone 74.5.2, Santa Cruz cat# sc-23885) or anti-PD-L1 (clone 10F.9G2, BioXcell cat# BE0101). The plate was incubated for 48 hours at 37°C and developed per manufacturer’s instructions.

#### Metabolic profiling

A single lung cell suspension was generated as above, followed by further enrichment of CD8^+^ T cells by magnetic bead separation (Miltenyi Biotec, cat# 130-104-075) per manufacturer’s instructions. CD8^+^ T cells were then counted using a BD Accuri^TM^ cytometer and plated with 200,000 cells/well with 10µg/mL anti-gC1qR. Metabolic status was assessed using Agilent Seahorse XF Cell Mito Stress Test Kit (cat# 103010-100) per manufacturer’s instructions on the Agilent Seahorse XFe96 analyzer.

#### C1q staining on autopsy specimens

Formalin fixed, paraffin embedded tissues were stained on the Ventana BenchMark Ultra automated staining platform. Slides were pretreated with ultraCC1 (Roche Tissue Diagnostics Indianapolis, IN) and stained using a polyclonal FITC labeled anti-C1q primary antibody (Roche Tissue Diagnostics, Indianapolis, IN, cat# 05267994001); sections were counterstained with DAPI. Slides were viewed using an Olympus BX63 microscope equipped with appropriate FITC and DAPI filter sets.

### Quantification and Statistical Analysis

All data displayed as mean±SEM. Analyses with two groups were analyzed using unpaired student’s t tests, while analyses with three or more groups were analyzed using a one-way ANOV test with multiple comparisons. Significance was defined as p<0.05 for all analyses. All statistical analysis was performed using GraphPad Prism (v.9.3.0).

### Key resources table

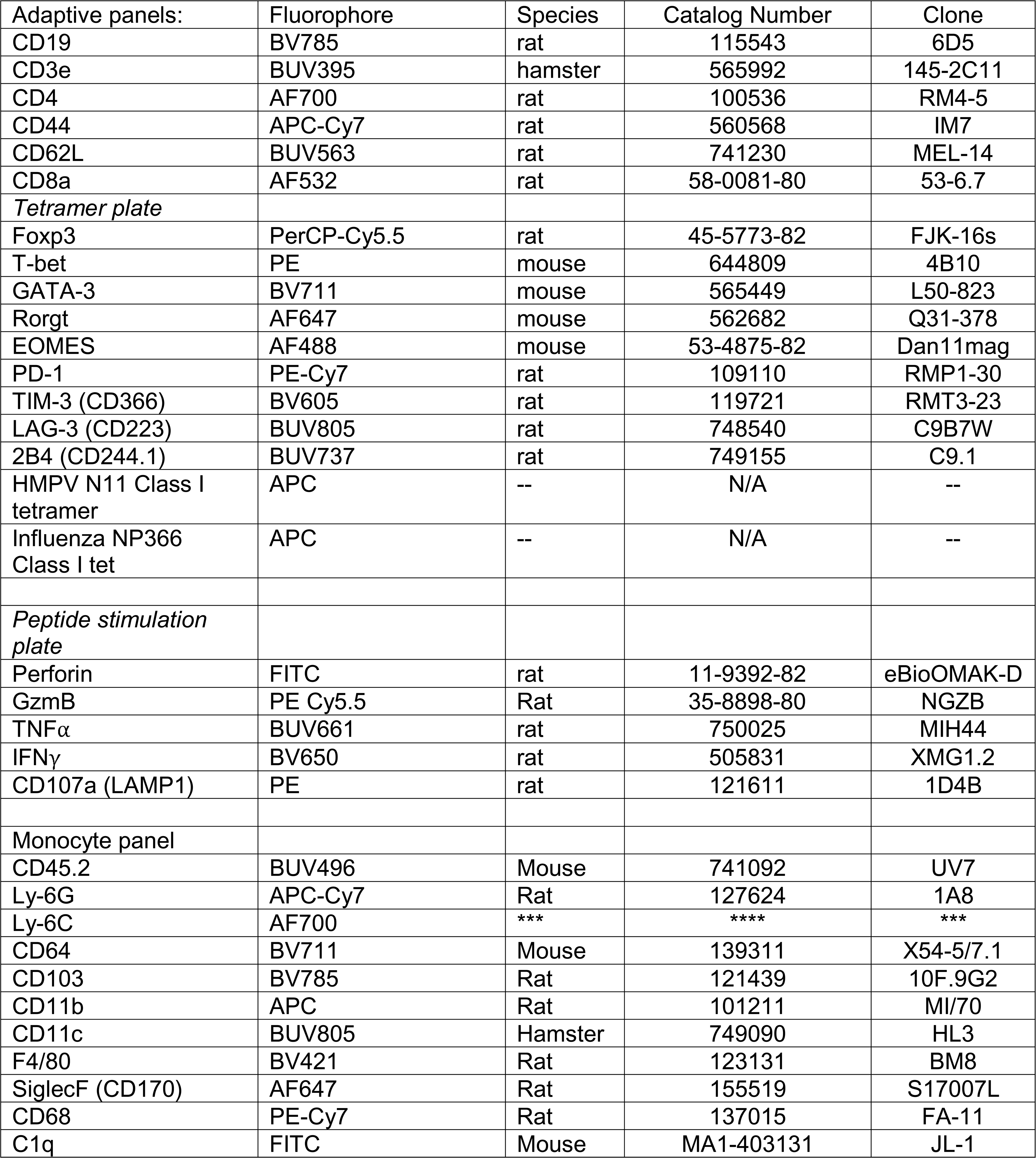

## References

Aegerter, H., Lambrecht, B.N., and Jakubzick, C.V. (2022). Biology of lung macrophages in health and disease. Immunity 55, 1564–1580.

Browaeys, R., Saelens, W., and Saeys, Y. (2020). NicheNet: modeling intercellular communication by linking ligands to target genes. Nat. Methods 17, 159–162.

Bulla, R., Tripodo, C., Rami, D., Ling, G.S., Agostinis, C., Guarnotta, C., Zorzet, S., Durigutto, P., Botto, M., and Tedesco, F. (2016). C1q acts in the tumour microenvironment as a cancer-promoting factor independently of complement activation. Nat. Commun. 7, 10346.

Chang, J., and Braciale, T.J. (2002). Respiratory syncytial virus infection suppresses lung CD8+ T-cell effector activity and peripheral CD8+ T-cell memory in the respiratory tract. Nat. Med. 8, 54–60.

Channappanavar, R., Fehr, A.R., Vijay, R., Mack, M., Zhao, J., Meyerholz, D.K., and Perlman, S. (2016). Dysregulated Type I Interferon and Inflammatory Monocyte-Macrophage Responses Cause Lethal Pneumonia in SARS-CoV-Infected Mice. Cell Host Microbe 19, 181–193.

Chen, A., Gaddipati, S., Hong, Y., Volkman, D.J., Peerschke, E.I., and Ghebrehiwet, B. (1994). Human T cells express specific binding sites for C1q. Role in T cell activation and proliferation. J. Immunol. 153, 1430–1440.

Coates, B.M., Staricha, K.L., Koch, C.M., Cheng, Y., Shumaker, D.K., Budinger, G.R.S., Perlman, H., Misharin, A.V., and Ridge, K.M. (2018). Inflammatory Monocytes Drive Influenza A Virus-Mediated Lung Injury in Juvenile Mice. J. Immunol. 200, 2391–2404.

Erickson, J.J., Gilchuk, P., Hastings, A.K., Tollefson, S.J., Johnson, M., Downing, M.B., Boyd, K.L., Johnson, J.E., Kim, A.S., Joyce, S., et al. (2012). Viral acute lower respiratory infections impair CD8+ T cells through PD-1. J. Clin. Invest. 122, 2967–2982.

Erickson, J.J., Rogers, M.C., Hastings, A.K., Tollefson, S.J., and Williams, J.V. (2014). Programmed death-1 impairs secondary effector lung CD8^+^ T cells during respiratory virus reinfection. J. Immunol. 193, 5108–5117.

Erickson, J.J., Rogers, M.C., Tollefson, S.J., Boyd, K.L., and Williams, J.V. (2016). Multiple Inhibitory Pathways Contribute to Lung CD8+ T Cell Impairment and Protect against Immunopathology during Acute Viral Respiratory Infection. J. Immunol. 197, 233–243.

Foley, D.A., Sikazwe, C.T., Minney-Smith, C.A., Ernst, T., Moore, H.C., Nicol, M.P., Smith, D.W., Levy, A., and Blyth, C.C. (2022). An Unusual Resurgence of Human Metapneumovirus in Western Australia Following the Reduction of Non-Pharmaceutical Interventions to Prevent SARS-CoV-2 Transmission. Viruses 14.

GBD 2019 LRI Collaborators (2022). Age-sex differences in the global burden of lower respiratory infections and risk factors, 1990-2019: results from the Global Burden of Disease Study 2019. Lancet Infect. Dis. 22, 1626–1647.

Ghebrehiwet, B., Geisbrecht, B.V., Xu, X., Savitt, A.G., and Peerschke, E.I.B. (2019). The C1q Receptors: Focus on gC1qR/p33 (C1qBP, p32, HABP-1)1. Semin. Immunol. 45, 101338.

Grant, R.A., Morales-Nebreda, L., Markov, N.S., Swaminathan, S., Querrey, M., Guzman, E.R., Abbott, D.A., Donnelly, H.K., Donayre, A., Goldberg, I.A., et al. (2021). Circuits between infected macrophages and T cells in SARS-CoV-2 pneumonia. Nature 590, 635–641.

Jin, S., Guerrero-Juarez, C.F., Zhang, L., Chang, I., Ramos, R., Kuan, C.-H., Myung, P., Plikus, M.V., and Nie, Q. (2021). Inference and analysis of cell-cell communication using CellChat. Nat. Commun. 12, 1088.

Kishore, U., and Reid, K.B. (2000). C1q: structure, function, and receptors. Immunopharmacology 49, 159–170.

Knoll, R., Schultze, J.L., and Schulte-Schrepping, J. (2021). Monocytes and Macrophages in COVID-19. Front. Immunol. 12, 720109.

Lawrence, C.W., Ream, R.M., and Braciale, T.J. (2005). Frequency, specificity, and sites of expansion of CD8+ T cells during primary pulmonary influenza virus infection. J. Immunol. 174, 5332–5340.

Leach, S.M., Gibbings, S.L., Tewari, A.D., Atif, S.M., Vestal, B., Danhorn, T., Janssen, W.J., Wager, T.D., and Jakubzick, C.V. (2020). Human and Mouse Transcriptome Profiling Identifies Cross-Species Homology in Pulmonary and Lymph Node Mononuclear Phagocytes. Cell Rep. 33, 108337.

Li, Y., and Nair, H. (2022). Trends in the global burden of lower respiratory infections: the knowns and the unknowns. Lancet Infect. Dis. 22, 1523–1525.

Li, X., Kolling, F.W., Aridgides, D., Mellinger, D., Ashare, A., and Jakubzick, C.V. (2022). ScRNA-seq expression of IFI27 and APOC2 identifies four alveolar macrophage superclusters in healthy BALF. Life Sci. Alliance 5.

Liao, M., Liu, Y., Yuan, J., Wen, Y., Xu, G., Zhao, J., Cheng, L., Li, J., Wang, X., Wang, F., et al. (2020). Single-cell landscape of bronchoalveolar immune cells in patients with COVID-19. Nat. Med. 26, 842–844.

Ling, G.S., Crawford, G., Buang, N., Bartok, I., Tian, K., Thielens, N.M., Bally, I., Harker, J.A., Ashton-Rickardt, P.G., Rutschmann, S., et al. (2018). C1q restrains autoimmunity and viral infection by regulating CD8+ T cell metabolism. Science 360, 558–563.

Markarian, M., Krattli, R.P., Baddour, J.D., Alikhani, L., Giedzinski, E., Usmani, M.T., Agrawal, A., Baulch, J.E., Tenner, A.J., and Acharya, M.M. (2021). Glia-Selective Deletion of Complement C1q Prevents Radiation-Induced Cognitive Deficits and Neuroinflammation. Cancer Res. 81, 1732–1744.

Martin, M.D., and Badovinac, V.P. (2018). Defining memory CD8 T cell. Front. Immunol. 9, 2692.

Melms, J.C., Biermann, J., Huang, H., Wang, Y., Nair, A., Tagore, S., Katsyv, I., Rendeiro, A.F., Amin, A.D., Schapiro, D., et al. (2021). A molecular single-cell lung atlas of lethal COVID-19. Nature 595, 114–119.

Merad, M., and Martin, J.C. (2020). Pathological inflammation in patients with COVID-19: a key role for monocytes and macrophages. Nat. Rev. Immunol. 20, 355–362.

Nuriev, R., and Johansson, C. (2019). Chemokine regulation of inflammation during respiratory syncytial virus infection. [version 1; peer review: 3 approved]. F1000Res. 8.

Ogawa, T., Shichino, S., Ueha, S., Bando, K., and Matsushima, K. (2022). Profibrotic properties of C1q+ interstitial macrophages in silica-induced pulmonary fibrosis in mice. Biochem. Biophys. Res. Commun. 599, 113–119.

Olsen, S.J., Winn, A.K., Budd, A.P., Prill, M.M., Steel, J., Midgley, C.M., Kniss, K., Burns, E., Rowe, T., Foust, A., et al. (2021). Changes in influenza and other respiratory virus activity during the COVID-19 pandemic-United States, 2020-2021. Am. J. Transplant. 21, 3481–3486.

Roumenina, L.T., Daugan, M.V., Noé, R., Petitprez, F., Vano, Y.A., Sanchez-Salas, R., Becht, E., Meilleroux, J., Clec’h, B.L., Giraldo, N.A., et al. (2019). Tumor Cells Hijack Macrophage-Produced Complement C1q to Promote Tumor Growth. Cancer Immunol Res 7, 1091–1105.

Ruckwardt, T.J., Bonaparte, K.L., Nason, M.C., and Graham, B.S. (2009). Regulatory T cells promote early influx of CD8+ T cells in the lungs of respiratory syncytial virus-infected mice and diminish immunodominance disparities. J. Virol. 83, 3019–3028.

Schmidt, M.E., and Varga, S.M. (2018). The CD8 T cell response to respiratory virus infections. Front. Immunol. 9, 678.

Singh, R., Hemati, H., Bajpai, M., Yadav, P., Maheshwari, A., Kumar, S., Agrawal, S., Sevak, J.K., Islam, M., Mars, J.S., et al. (2021). Sustained expression of inflammatory monocytes and activated T cells in COVID-19 patients and recovered convalescent plasma donors. Immun. Inflamm. Dis. 9, 1279–1290.

Son, M. (2022). Understanding the contextual functions of C1q and LAIR-1 and their applications. Exp Mol Med 54, 567–572.

Sun, J., Madan, R., Karp, C.L., and Braciale, T.J. (2009). Effector T cells control lung inflammation during acute influenza virus infection by producing IL-10. Nat. Med. 15, 277–284.

Thielens, N.M., Tedesco, F., Bohlson, S.S., Gaboriaud, C., and Tenner, A.J. (2017). C1q: A fresh look upon an old molecule. Mol. Immunol. 89, 73–83.

Tian, H., Wang, G., Wang, Q., Zhang, B., Jiang, G., Li, H., Chai, D., Fang, L., Wang, M., and Zheng, J. (2022). Complement C1q binding protein regulates T cells’ mitochondrial fitness to affect their survival, proliferation, and anti-tumor immune function. Cancer Sci. 113, 875–890.

Williams, J.V., Tollefson, S.J., Johnson, J.E., and Crowe, J.E. (2005). The cotton rat (Sigmodon hispidus) is a permissive small animal model of human metapneumovirus infection, pathogenesis, and protective immunity. J. Virol. 79, 10944–10951.

Xu, J., Zhang, Y., and Williams, J.V. (2018). Development and optimization of a direct plaque assay for trypsin-dependent human metapneumovirus strains. J. Virol. Methods 259, 1–9.

Yao, Z.Q., Eisen-Vandervelde, A., Waggoner, S.N., Cale, E.M., and Hahn, Y.S. (2004). Direct binding of hepatitis C virus core to gC1qR on CD4+ and CD8+ T cells leads to impaired activation of Lck and Akt. J. Virol. 78, 6409–6419.

Zhou, Y., Fu, B., Zheng, X., Wang, D., Zhao, C., Qi, Y., Sun, R., Tian, Z., Xu, X., and Wei, H. (2020). Pathogenic T-cells and inflammatory monocytes incite inflammatory storms in severe COVID-19 patients. Natl Sci Rev 7, 998–1002.

