## Supplemental Figures for "Monocyte production of C1q potentiates CD8^+^ T cell effector function following respiratory viral infection"

† denotes co-first authorship

*Corresponding author. **Corresponding Author:**

John V. Williams, MD

Supplemental Figures 1-8

Supplemental Table 1

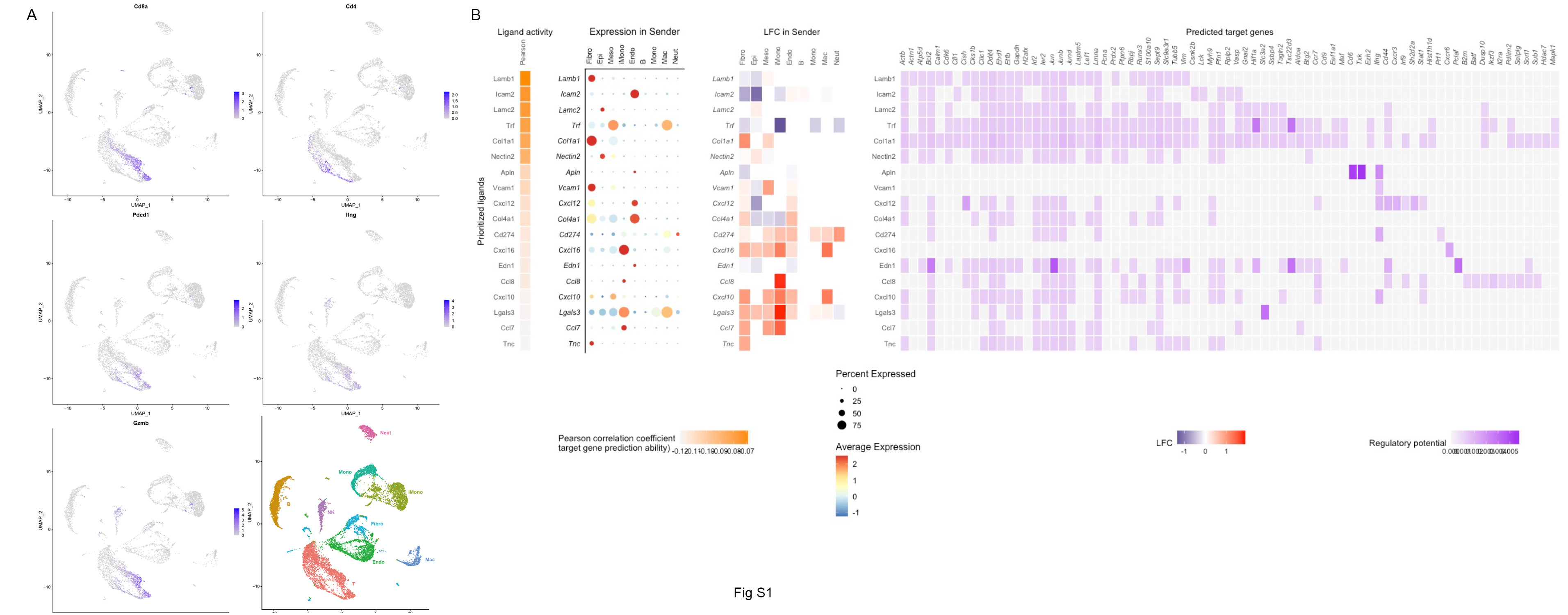

**Supplemental Figure 1**. **Validation of CD8 effector function and inhibitory receptor expression by single cell RNA sequencing.** A.) UMAPs of *Cd8a, Cd4, Pdcd1* (protein: PD-1), *Ifng*, and *Gzmb* showing co-expression of effector functions and PD-1 in CD8s. B.) NicheNet analysis of upregulated ligands (left), including PD-L1 (gene: *Cd274*), and the relative expression in sender populations (middle). Regulation of target genes (right) by individual ligands reveals PD-L1 regulation of *Ifng*.

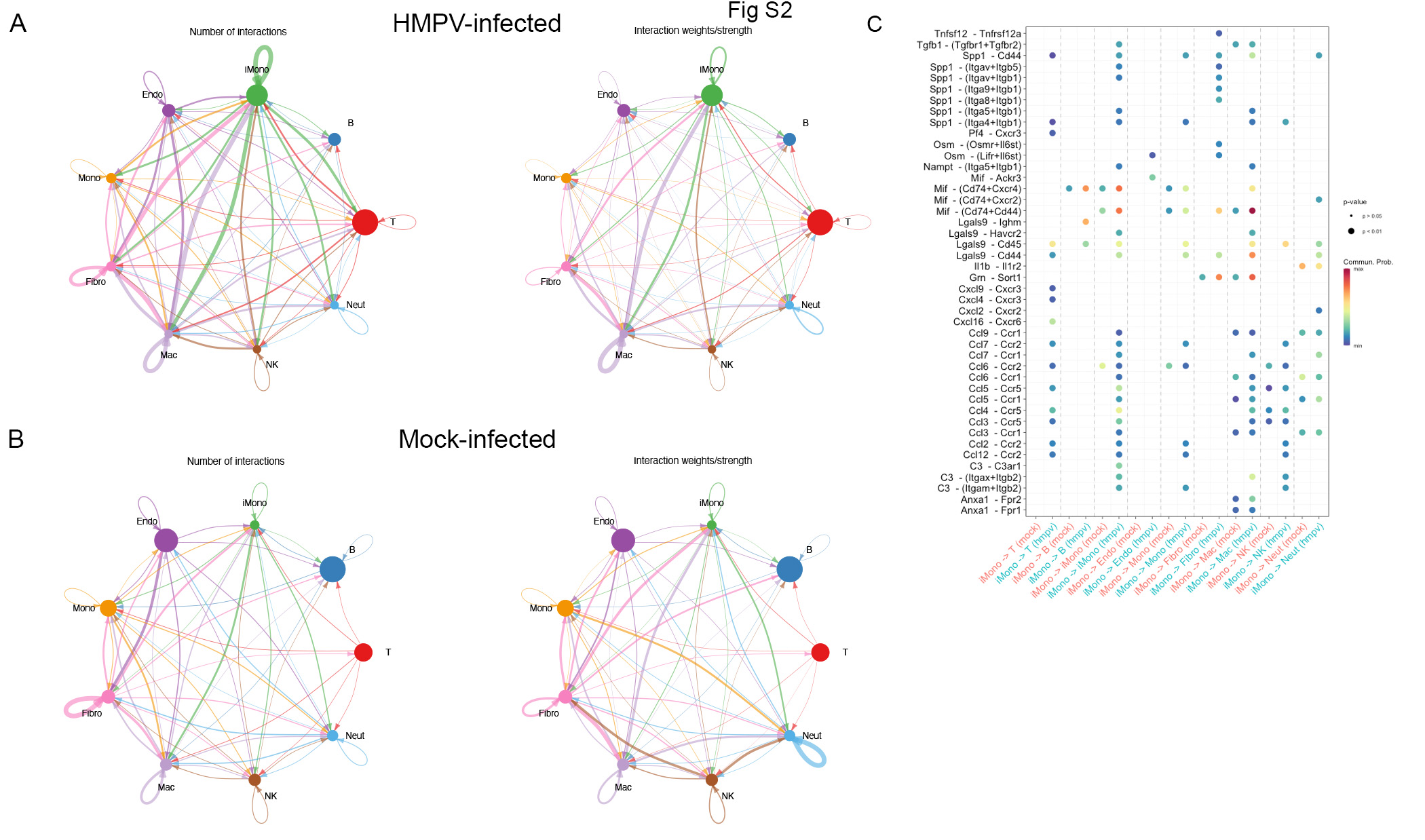

**Supplemental Figure 2**. **Cell-to-cell communication networks following HMPV-infection.** A-B.) Number of interactions and interaction strength between 9 cell populations following HMPV infection and mock infection, showing iMono-dominant communication after infection. C.) iMono signals transmitted to other cell populations in mock infection (red) or HMPV-infection (blue).

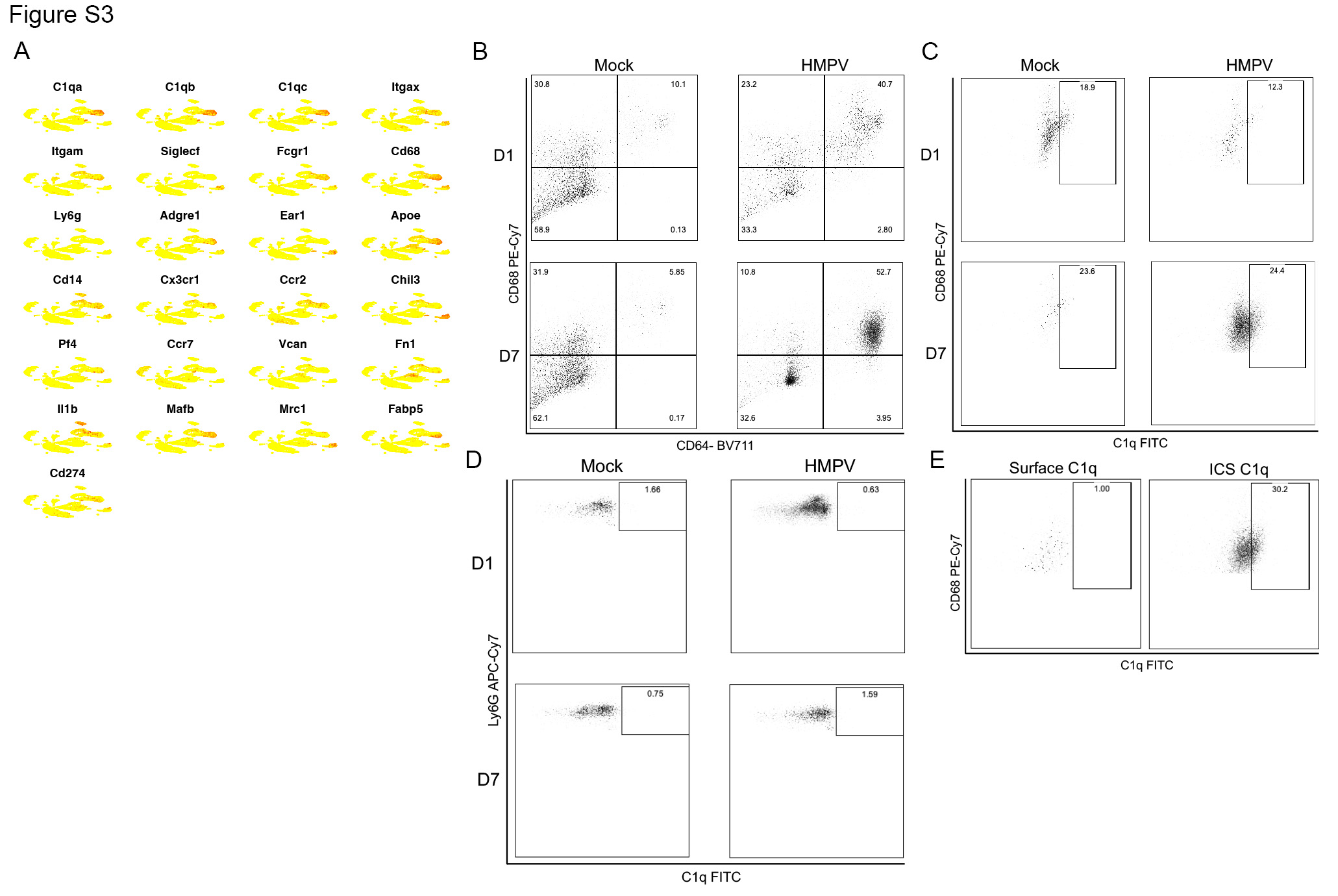

**Supplemental Figure 3**. **C1q production by inflammatory monocytes on day 7 after HMPV infection**. A.) Expression of several monocyte/macrophage gene transcripts overlaid on a UMAP, showing upregulation of CD64 (gene: *Fcgr1*) and *Cd68* in C1qa-producing population. B.) CD64^+^CD68^+^ cell recruitment on day 7 after HMPV infection. C.) C1q production in CD64^+^CD68^+^ cells by intracellular staining. D.) Similar staining on Ly6G^+^ neutrophils. E.) Limited C1q staining without the use of permeabilization (left) compared to intracellular staining (ICS) on CD64^+^CD68^+^ cells.

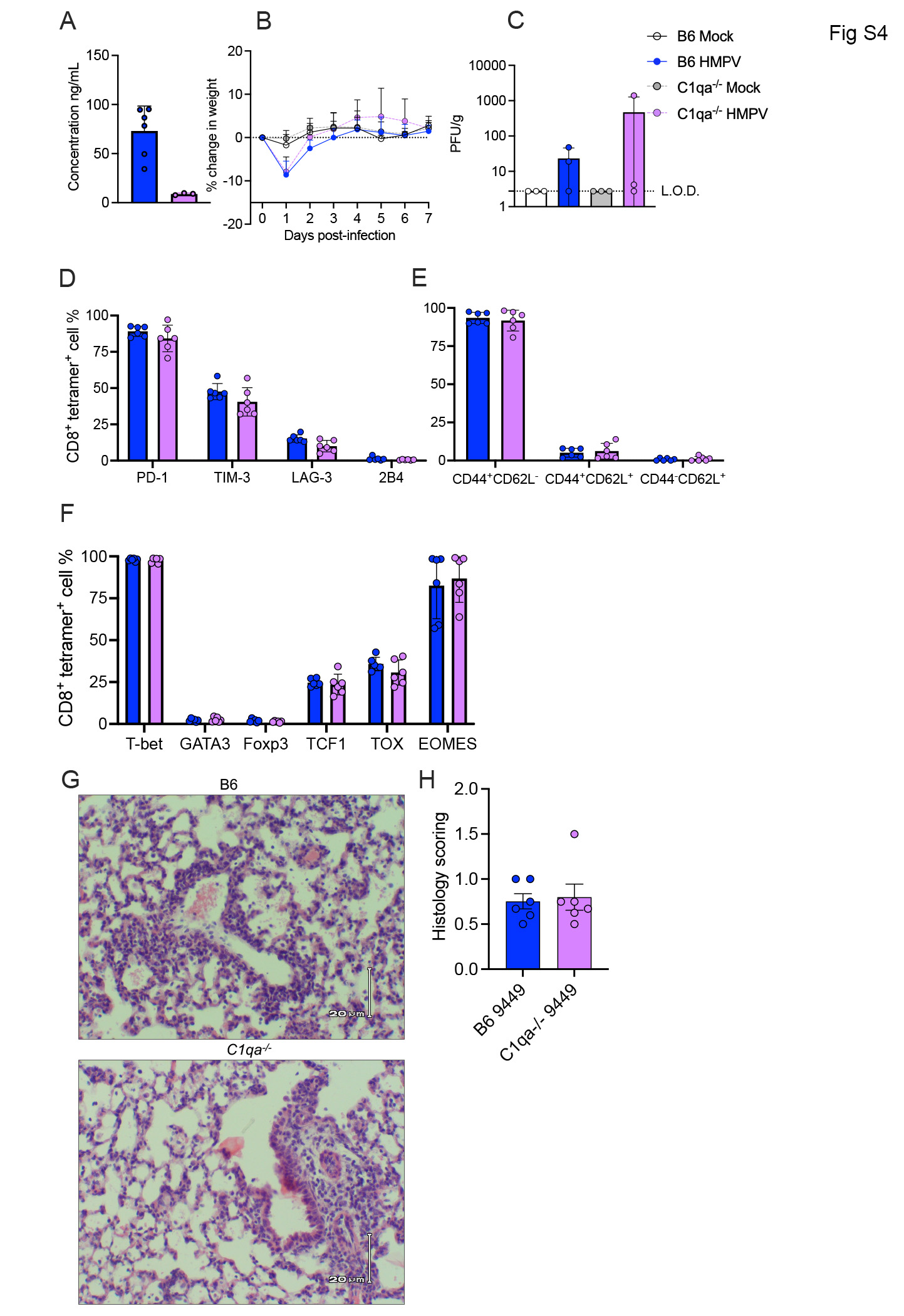

**Supplemental Figure 4**. **Inhibitory receptor, activation states, and transcription factor expression in B6 vs *C1qa^-/-^* mice.** A.) C1q quantity in B6 vs. *C1qa^-/-^* mice. B.) No changes in weight between groups. C.) Low burdens of HMPV infection in infected animals regardless of genotype. D.) Expression of PD-1, TIM-3, LAG-3, and 2B4 in B6 and *C1qa^-/-^* HMPV-infected mice. E.) CD44/CD62L activation states in in B6 and *C1qa^-/-^* HMPV-infected mice. F.) Expression of transcription factors in in B6 and *C1qa^-/-^* HMPV-infected mice. G-H.) Representative histopathology staining and quantification at day 7 post infection in in B6 and *C1qa^-/-^* HMPV-infected mice.

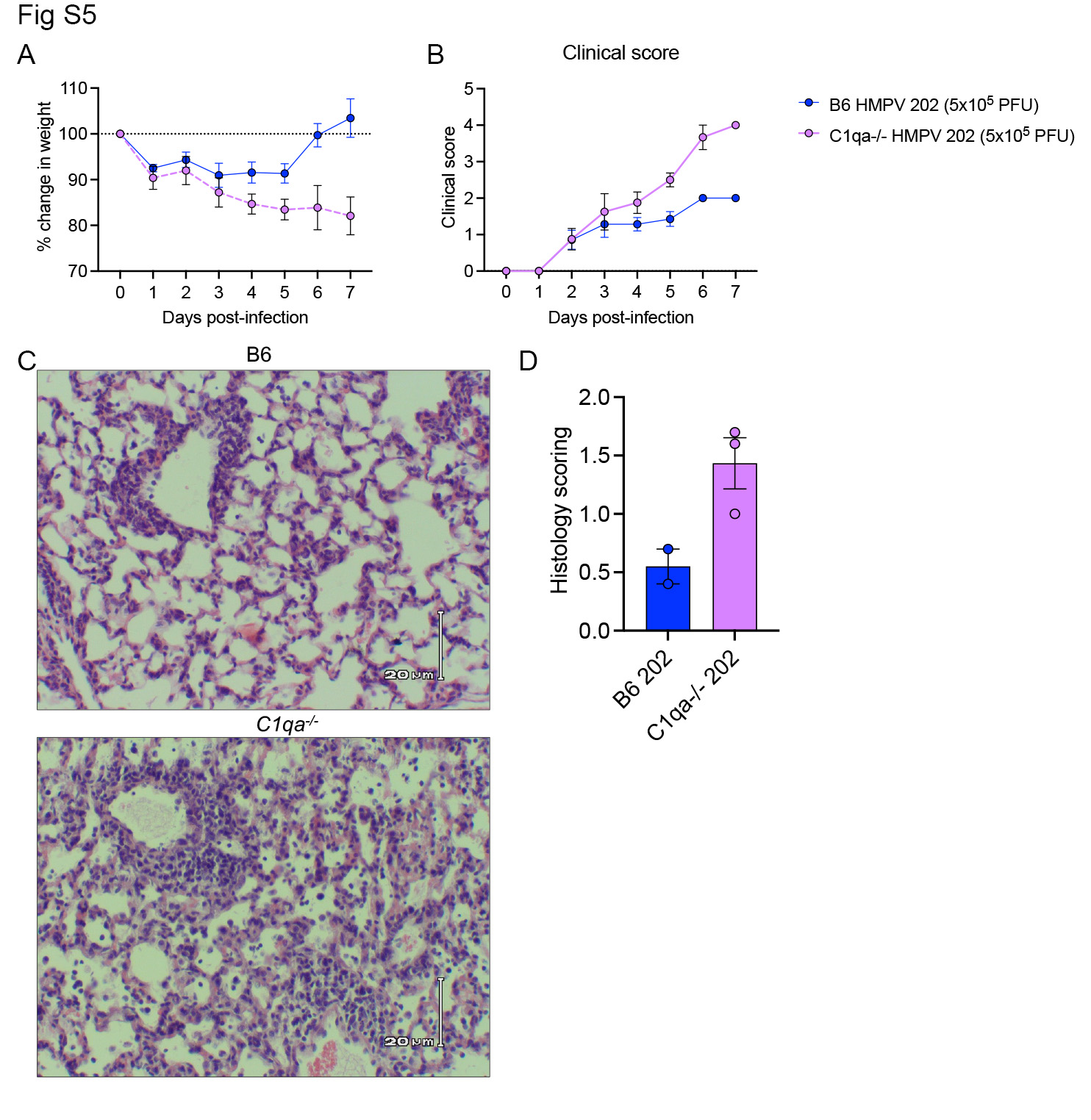

**Supplemental Figure 5**. **Virulent HMPV strain leads to enhanced disease in *C1qa^-/-^* mice.** A.) Increased weight loss in *C1qa^-/-^ mice* infected with C-202. B.) Increased clinical score in *C1qa^-/-^ mice* infected with C-202 calculated by hunched, fur grooming, respiratory rate, and activity. C.) Representative histopathology staining and quantification at day 7 post infection in in B6 and *C1qa^-/-^* C-202-infected mice.

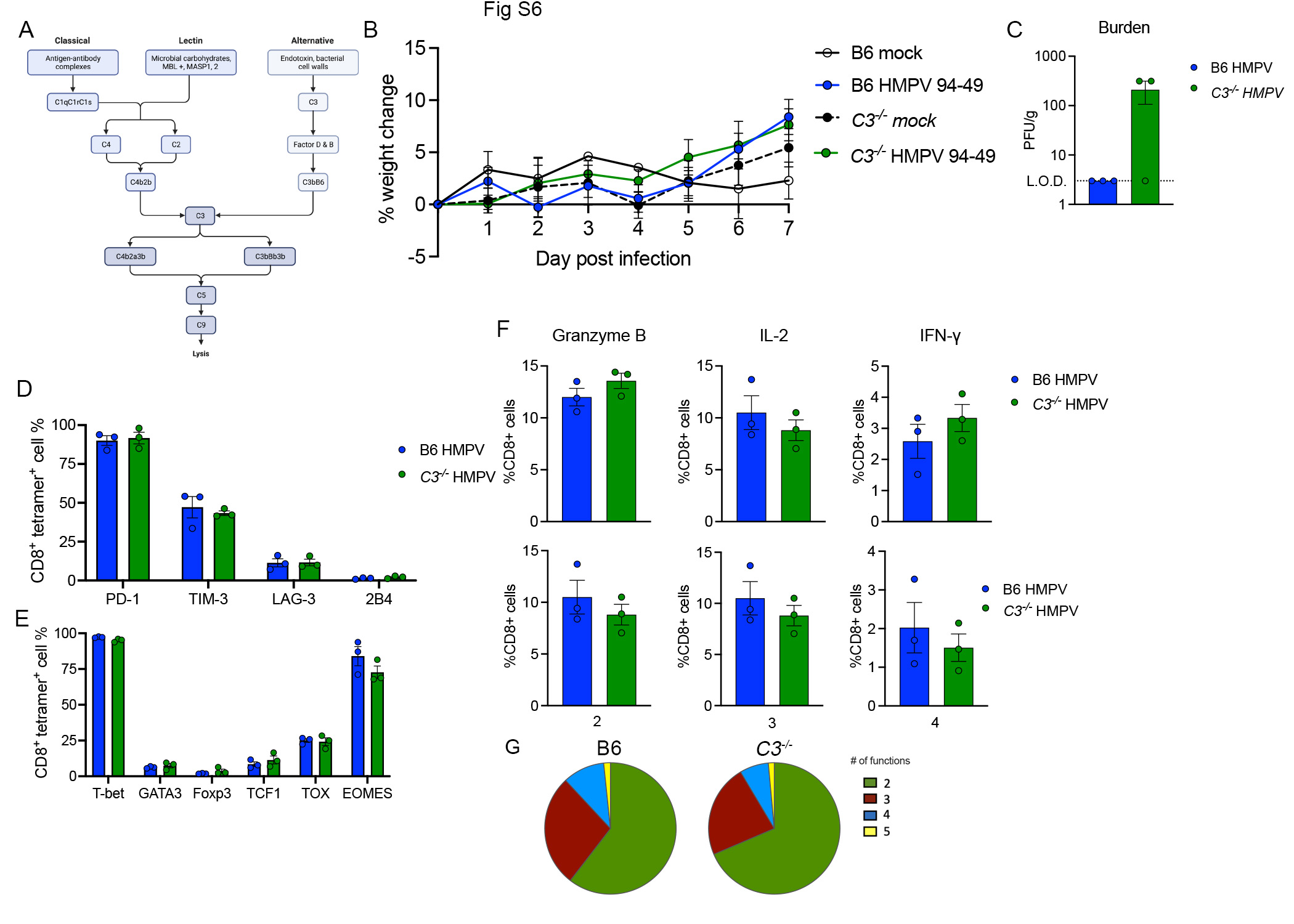

**Supplemental Figure 6**. **CD8 effector function is preserved in animals lacking C3**. A.) Schematic of complement pathways showing convergence on C3 (Biorender). B.) No changes in weight between groups. C.) Low burdens of HMPV infection in infected animals regardless of genotype. D.) Expression of PD-1, TIM-3, LAG-3, and 2B4 in B6 and *C3^-/-^* HMPV-infected mice. E.) Expression of transcription factors in B6 and *C3^-/-^* HMPV-infected mice. F.) No changes in effector functions (granzyme B, IL-2, and IFN-γ) in B6 and *C3^-/-^* HMPV-infected mice. G.) Boolean gating demonstrating similar distribution of multiple effector functions in B6 and *C3^-/-^* HMPV-infected mice.

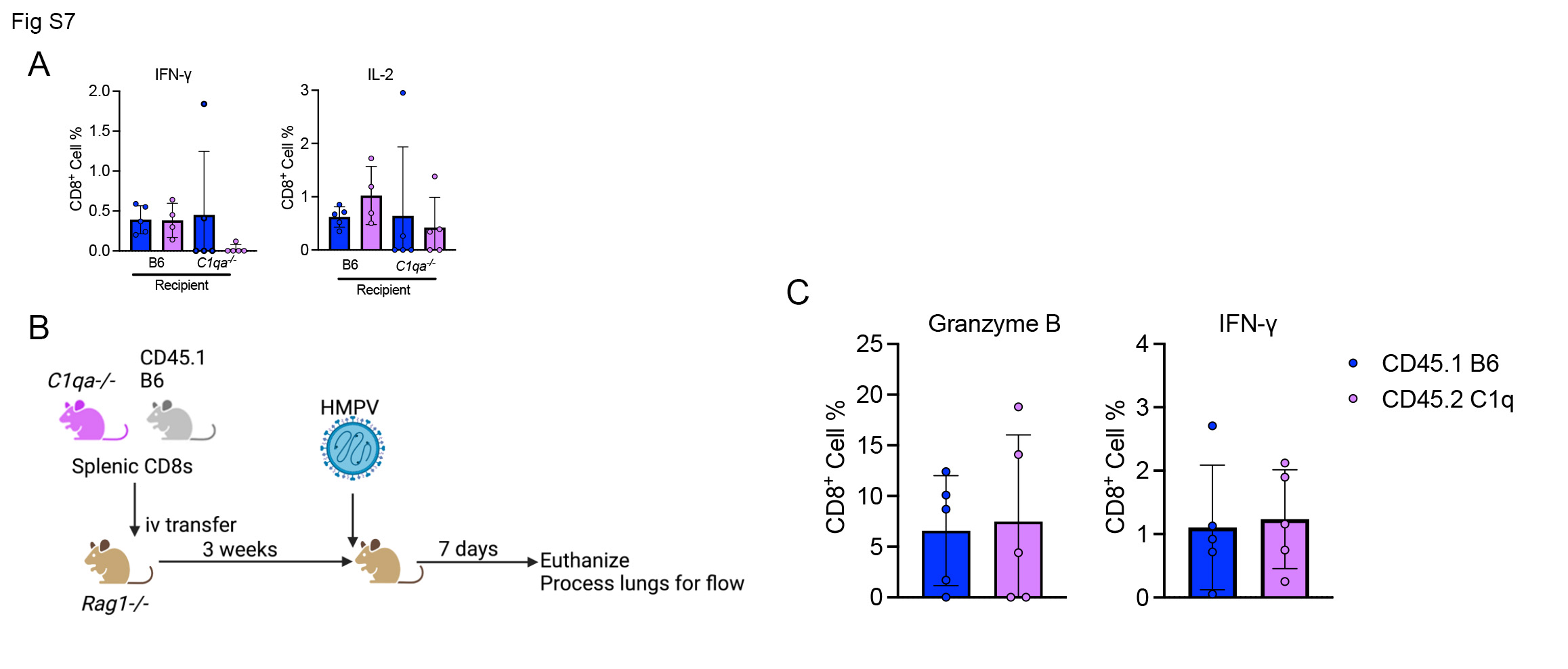

**Supplemental Figure 7**. **CD8 extrinsic requirement of C1q*.*** A.) Four-way adoptive transfer experiments showing no difference in IL-2 or IFN-γ. B.) Adoptive transfer schematic of either CD8s from B6 or *C1qa^-/-^* mice into *Rag1^-/-^* recipients followed by HMPV infection. C.) No change in granzyme B or IFN-γ production by B6 or *C1qa^-/-^* CD8s.

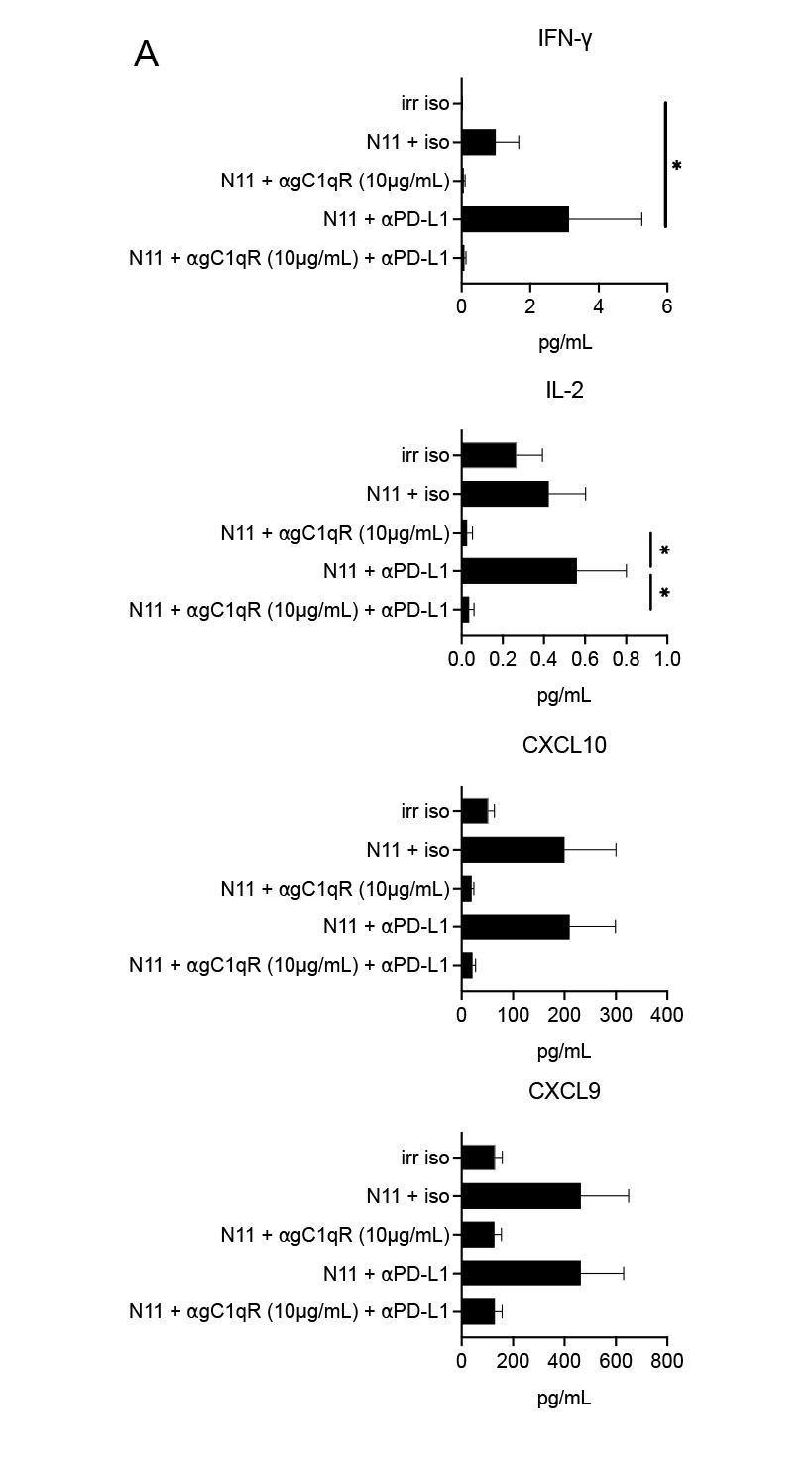

**Supplemental Figure 8. ⍺gC1qR treatment abrogates cytokine production regardless of PD-L1 blockade**. A.) IFN-γ, IL-2, CXCL10, and CXCL9 production following 48 hours of *ex vivo* stimulation with reduced cytokine production when treated with ⍺gC1qR. *p<0.05 by one-way ANOVA with multiple comparisons.

**Supplemental Table 1**: Key reagents.

| Adaptive panels: | Fluorophore | Species | Catalog Number | Clone |
| --- | --- | --- | --- | --- |
| CD19 | BV785 | rat | 115543 | 6D5 |
| CD3e | BUV395 | hamster | 565992 | 145-2C11 |
| CD4 | AF700 | rat | 100536 | RM4-5 |
| CD44 | APC-Cy7 | rat | 560568 | IM7 |
| CD62L | BUV563 | rat | 741230 | MEL-14 |
| CD8a | AF532 | rat | 58-0081-80 | 53-6.7 |
| *Tetramer plate* |  |  |  |  |
| Foxp3 | PerCP-Cy5.5 | rat | 45-5773-82 | FJK-16s |
| T-bet | PE | mouse | 644809 | 4B10 |
| GATA-3 | BV711 | mouse | 565449 | L50-823 |
| Rorgt | AF647 | mouse | 562682 | Q31-378 |
| EOMES | AF488 | mouse | 53-4875-82 | Dan11mag |
| PD-1 | PE-Cy7 | rat | 109110 | RMP1-30 |
| TIM-3 (CD366) | BV605 | rat | 119721 | RMT3-23 |
| LAG-3 (CD223) | BUV805 | rat | 748540 | C9B7W |
| 2B4 (CD244.1) | BUV737 | rat | 749155 | C9.1 |
| HMPV N11 Class I tetramer | APC | -- | N/A | -- |
| Influenza NP366 Class I tet | APC | -- | N/A | -- |
| *Peptide stimulation plate* |  |  |  |  |
| Perforin | FITC | rat | 11-9392-82 | eBioOMAK-D |
| GzmB | PE Cy5.5 | Rat | 35-8898-80 | NGZB |
| TNF$\alpha$ | BUV661 | rat | 750025 | MIH44 |
| IFN$\gamma$ | BV650 | rat | 505831 | XMG1.2 |
| CD107a (LAMP1) | PE | rat | 121611 | 1D4B |
| Monocyte panel |  |  |  |  |
| CD45.2 | BUV496 | Mouse | 741092 | UV7 |
| Ly-6G | APC-Cy7 | Rat | 127624 | 1A8 |
| Ly-6C | AF700 | *** | **** | *** |
| CD64 | BV711 | Mouse | 139311 | X54-5/7.1 |
| CD103 | BV785 | Rat | 121439 | 10F.9G2 |
| CD11b | APC | Rat | 101211 | MI/70 |
| CD11c | BUV805 | Hamster | 749090 | HL3 |
| F4/80 | BV421 | Rat | 123131 | BM8 |
| SiglecF (CD170) | AF647 | Rat | 155519 | S17007L |
| CD68 | PE-Cy7 | Rat | 137015 | FA-11 |
| C1q | FITC | Mouse | MA1-403131 | JL-1 |
